# DNA extracted from boiled archival fish bones yields high quality whole genome sequencing data

**DOI:** 10.1101/2025.06.10.658604

**Authors:** J Niu, A Vasemägi, M-E López, L Pukk, M Huss, A Gårdmark

## Abstract

Archival samples provide a unique source of DNA, offering the potential to extend the temporal scale of genetic studies by decades to centuries. Fish hard structures, such as otoliths and scales, serve as records for fish collected during fisheries monitoring across a large spatiotemporal scale. Operculum bones are a type of fish hard structure that, although less commonly collected, have been utilized in genetic studies. However, the usefulness of archived operculum bones to provide high quality whole genome sequence information has not been thoroughly evaluated. Here, we applied a commercially available extraction protocol, with minor adjustments, to isolate genomic DNA from operculum bones of Eurasian perch collected up to 47 years ago, followed by standard short-read whole genome sequencing. By comparing the metrics of DNA and whole genome sequencing data of bones with contemporary muscle samples, we demonstrate that the application of a simple extraction protocol on boiled archival fish operculum bones yields high quality genomic information that is comparable to fresh tissue, highlighting the role of archival operculum bones as an overlooked repository of valuable genomic information.

## INTRODUCTION

DNA analyses from historical samples provide opportunities to elucidate how past events or environmental conditions have shaped demographic history of populations and species, track evolutionary changes over time, and to directly measure the rate of molecular evolution. Since the 1980s, researchers have studied ancient DNA (aDNA) stored in archaeological remains of humans, domestic animals and extinct megafauna (Pääbo 1993; Hofreiter et al. 2015). In aquatic organisms, genetic data has been successfully sequenced from DNA isolated from fish bones stemming from the late Holocene (Ferrari et al. 2021; Martínez-García et al. 2022). Skeletal remains may actually surpass soft tissue as a repository for DNA, potentially due to DNA’s affinity to bind to bone minerals (Latham and Miller 2018). However, analyses of aDNA have proven challenging, both in terms of acquiring biological material in archaeological excavations (Rizzi et al. 2012), and because of the need for rigorous decontamination and quality control processes than can require significant time, effort, and expense (Gilbert et al. 2006). Even when such decontamination regimes and authenticity protocols are followed, contaminants are frequently observed (Kolman and Tuross 2000). Furthermore, the opportunistic nature of most archaeological material often limits their temporal resolution, complicating efforts to associate genetic finding with climate induced temperature changes and other anthropogenic stressors (Atmore et al. 2023; Ferrari et al. 2021).

Here, we focus on archival samples - another category of historical samples from which a significantly larger number of specimens have been consistently and regularly collected over the past 200 years (Wandeler, Hoeck, and Keller 2007). Due to the extensive efforts of fisheries institutes worldwide and their routine monitoring practices, there is a substantial collection of fish bony archival samples, including otoliths, scales, operculum bones, vertebrae, jaws, and other parts of the bony endoskeleton (Tzadik et al. 2017; Nielsen and Hansen 2008; Campana and Thorrold 2001). A significant portion of these bony samples was collected for determining fish age and growth, as they display visible annuli reflecting fish growth (Lai et al. 1995; Maunder and Punt 2013). While age determination is typically performed using otoliths and scales, the operculum bone, located on the surface of the fish head as part of the gill cover (see Figure 1 for an example in Eurasian perch), serves as a better alternative for age determination for some fish species (Le Cren 1947; McConell 1951) and for morphological or chemical analyses due to its flat shape and relatively large size (Huycke, Eames, and Kimmel 2012; Kusznierz et al. 2023; Tarasco et al. 2017). As a result, large collections of operculum bones exist for many species (Thoresson 1996; Ma et al. 2011; Khan and Khan 2009; Perry and Casselman 2012).

**Figure 1.**
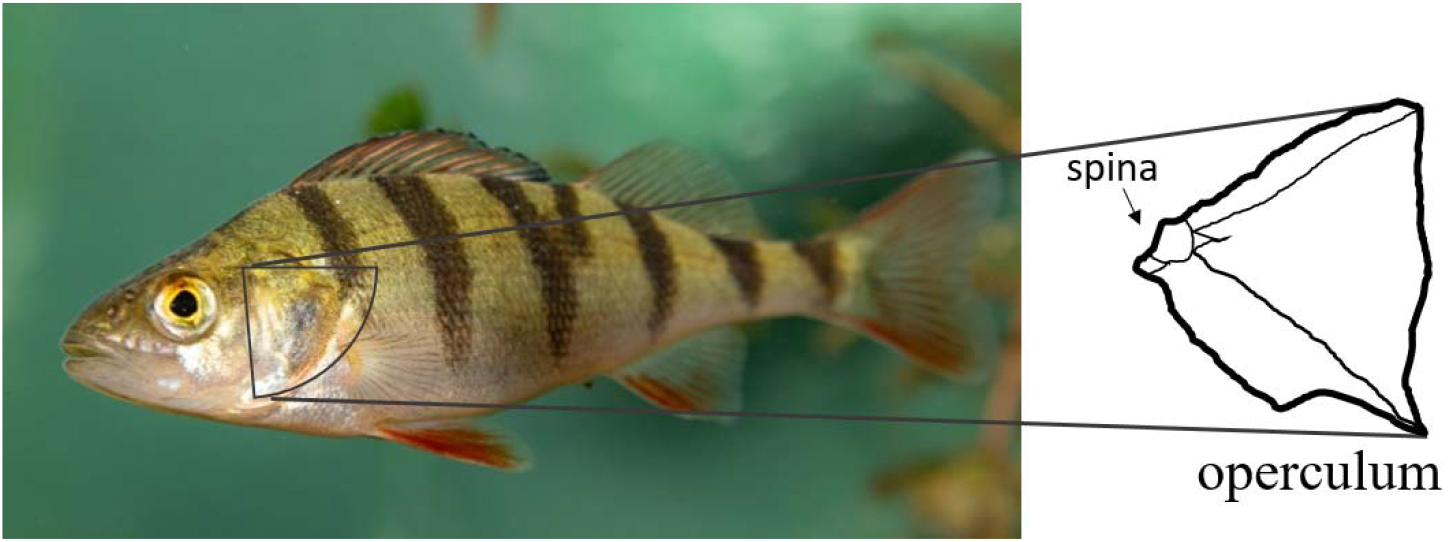
Schematic illustration of the operculum bone, which is part of the gill cover (here shown in a small Eurasian perch) and was used in DNA extraction. Adjusted photo of a Eurasian perch from Mark Harris.

Often stored in museums and research institutes, archives of fish bony samples constitute unparalleled spatiotemporal records of fish species and populations (Campana and Thorrold 2001; Nielsen and Hansen 2008). Since the late 1990s, the utility of archival samples as genetic repositories has been primarily used for investigating the temporal stability of population structure, genetic diversity and effective population size (Manuzzi et al. 2022; Swatdipong et al. 2010). Many studies have since identified genetic consequences from fishing on wild fish populations (Hutchinson et al. 2003; Pukk et al. 2013; Miettinen et al. 2024; Cuveliers et al. 2009; Price et al. 2019). Other concurrent stressors in the environment such as climate change, food availability, parasitic load, habitat connectivity have also been associated with evolutionary changes in fish through archival genetics (Czorlich et al. 2022; Bonanomi et al. 2015; Therkildsen et al. 2019; Björklund, Aho, and Behrmann-Godel 2015; McDermid et al. 2014). By analysing the genetic information collected from native, introduced and farmed populations, the genetic impact of historical stocking event has been revealed (Finnegan and Stevens 2008). Future stocking practice and spatial management units can be better informed and defined accordingly (Östergren et al. 2021; Ciborowski et al. 2007). Archival fish bony samples have demonstrated the ability to expand the time and space dimensions of evolutionary studies (Wandeler, Hoeck, and Keller 2007; Bi et al. 2013).

However, the majority of the molecular analyses performed using fish bony samples have been limited to a few mitochondrial DNA loci (Ciborowski et al. 2007) or microsatellite loci (Cuveliers et al. 2009; Price et al. 2019; Pukk et al. 2013). There is a growing number of studies utilising single nucleotide polymorphism (SNP) genotyping through reduced representation methods, such as restriction-site associated DNA sequencing (RADseq, Jacobsen et al. 2017), SNP chips (Johnston et al. 2013; Bonanomi et al. 2015) or genotyping-in-thousands by sequencing (GT-seq) (Setzke, Wong, and Russello 2021; Manuzzi et al. 2022). To our best knowledge, only studies that have generated whole-genome sequencing (WGS) data using archival bony samples from fish, to enable more robust evolutionary inferences and analyses of genomic variants beyond SNPs, have been limited to utilising fish scales and otoliths (Caccavo et al. 2024; Pinsky et al. 2021). Specifically, until now, no study has been done on archival operculum bones to test if such cellular bone tissue can yield DNA of enough quality to support subsequent whole-genome sequencing. Operculum bones may be particularly challenging for extraction of high-quality DNA as they – in contrast to otoliths and scales – are commonly treated with boiling water to remove skin and clean the bone.

In this study, we tested the possibility of generating whole-genome sequencing data by applying a commercially available DNA isolation kit, originally designed for blood and tissue samples, with only minor adjustments on archival operculum bones. Through a comparative analysis of DNA and sequencing quality from bones dated to the 1980s and 2000s, alongside fresh samples, we demonstrated that even boiled archival bones can effectively provide significant quantities of fish endogenous DNA in a manner that is both time-efficient and cost-effective, adequate for generating high-quality WGS data.

## MATERIAL AND METHODS

### The boiled operculum bones

Like many institutes working with fish and fisheries, the Department of Aquatic Resources at the Swedish University of Agricultural Sciences (SLU Aqua), holds archives of dried fish bony tissues, some of which date back to more than 100 years ago. Among these archives, there is one for operculum bones from Eurasian perch (*Perca fluviatilis*) that has been maintained since 1970. The archive has been established as an outcome of a coastal fish monitoring program that uses gillnets and fyke nets to track fish populations, and environmental changes over time (Sandström, Neuman, and Thoresson 1995; Adill et al. 2013; Thoresson 1996). The operculum bones were sampled and prepared as follows: 1) removal of the left operculum of each individual; 2) placing each operculum bone in a separate plastic chamber; 3) pouring boiling water to each chamber to cover the bone; 4) once the boiling water has cooled down, removal of attached soft tissue on the bone by brushing and rinsing with tap water, and 5) leaving the bone to dry in a small paper envelop before translocating them into an archive room. Thus, the archival bones have been treated using boiling water to deliberately remove attached tissues rich in DNA, which is far from optimal for isolating and preserving high quality DNA.

To evaluate (1) whether it is possible to extract DNA of enough quantity and quality from boiled archival operculum bones for whole genome sequencing and (2) whether the storage time of bones affects the quality of DNA and WGS data, we selected a subset of 173 bone samples from this archive (Table S1). The subset contained bones collected from 1977-1978 (hereafter 1980s bones) and 2001-2002 (2000s bones) from two perch populations on the Baltic Sea coast of Sweden. The bones were selected by size to achieve sufficient sample weight (> 10 mg). The body length of fish from which the operculum bones were sampled ranged from 166 to 340 mm (Figure S1).

### Contemporary samples

To further evaluate how the type of tissue (bone/muscle), boiling and storage time of the bones affect quality of DNA and WGS data, we also used 105 contemporary perch muscle tissue samples collected from gillnet-fishing in 2021 and 2022 from the same two populations. Approximately 1-3 cm^3^ of muscle was taken along the dorsal fin after removing the skin, using a clean scalpel and tweezers. The muscle samples were stored in 5 ml Eppendorf tubes filled with 95% ethanol and kept at 4°C. The tools and cutting board were cleaned and sterilized using 70% ethanol in between processing each individual. For simplicity, we refer to these muscle samples as the 2020s muscles.

### DNA extraction

All DNA extractions were conducted in a specifically designed DNA laboratory with as many steps as possible done under a stamina flow hood to minimize potential cross-contamination and a strict unidirectional movement to avoid contact between pre- and post-PCR samples. Disposable equipment, such as gloves, were changed after every potential contact with any sample (including touching the opening of any tubes). Non-disposable equipment such as tweezers and bead-beating beads and work surfaces were cleaned with Milli-Q water, bleach (0.5% sodium hypochlorite solution), and 70% ethanol. One extraction negative was included for every 24 samples.

For the 173 bones, the operculum bones were broken into smaller pieces to fit into 2 ml polypropylene screw cap micro tubes. To avoid contamination that can be introduced in additional handling of the bone pieces, we did not measure bone pieces weight used for each sample. Metal tweezers were used to break the bone if it was too hard to break by hand. One piece of every sample was re-archived into the original paper envelop for potential future studies. Then five 3 mm Tungsten Carbide Beads (QIAGEN) were added to the sample and homogenized by a PowerLyzer 24 Homogenizer (QIAGEN) for 10 cycles of 30 s at speed of ∼5000 m/s plus 30 s dwell time to avoid overheating. This resulted in a total of 10 minutes of homogenizing at the maximum speed at which the micro tubes did not start to crack. The homogenizing pulverized the bone samples. Subsequently, samples were centrifuged for 5 min at high speed (11000 g) to settle the bone powder.

For subsequent extraction steps, we chose the Macherey-Nagel NucleoSpin Tissue kit (Macherey-Nagel NucleoSpin Tissue kit #740952.250) and followed a slightly modified version of their standard protocol. The DNA extraction steps are summarized in Figure 2.

**Figure 2.**
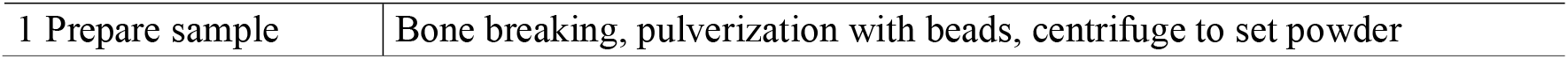
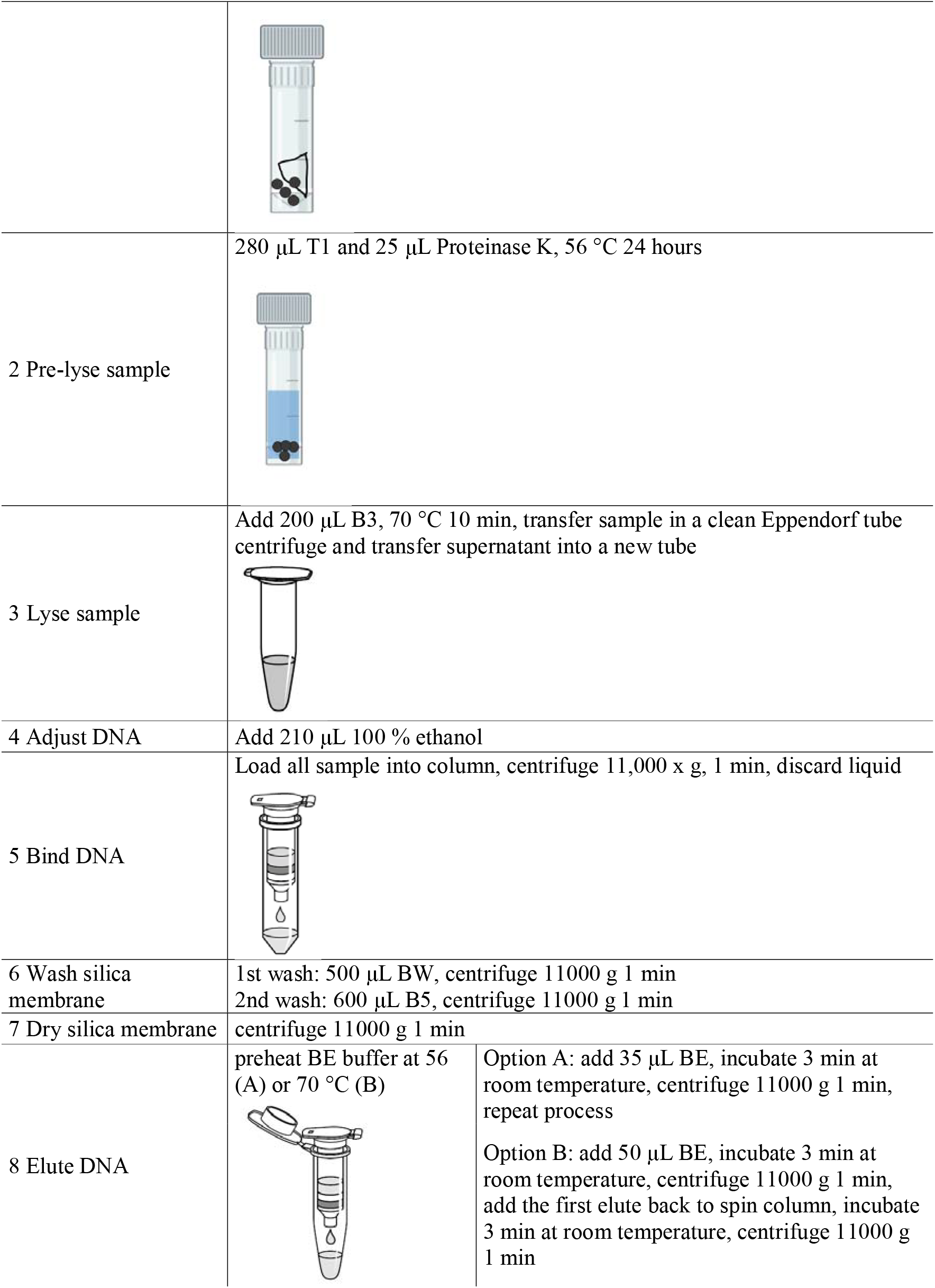
Graphic summary of the main steps of the DNA extraction protocol from fish operculum bones. Adapted from the Macherey-Nagel NucleoSpin Tissue kit manual.

For bone powder digestion, we increased the amount of reagents used at multiple steps. First, all samples were digested in 280 μl Buffer T1 and 25 μl of Proteinase K at 56°C for 24 hours. Tubes were vortexed twice during the process with even time intervals. Digestion was finished with the addition of 200 μl B3 Buffer and then kept at 70 °C for 10 minutes. To remove the Tungsten Carbide Beads from each sample, we transferred the supernatant from the micro tube to a sterile 1.5 ml Eppendorf tube. We centrifuged the 1.5 ml tube for 5 min at high speed (11000 g) and transferred the supernatant again to a new 1.5 ml Eppendorf tube, to remove any indigestible bone pieces from the sample before we proceeded with the DNA extraction.

We thereafter added 210 μl 100% ethanol to the sample and vortexed the mixture vigorously. The mixture was then loaded into the NucleoSpin Tissue Column. DNA was bound to the silica membrane inside the NucleoSpin Tissue Column during the process of the mixture being washed through during centrifuging (11000 g). We washed impurities off the silica membrane by using 500 μl Buffer BW and 600 μl Buffer B5 following the standard protocol.

Finally, to collect as much DNA bound to the silica membrane as possible, we experimented with two alternative elution steps (A and B) on different bone samples to maximize the elution efficiency (Table S1). Option A using 56 °C Elution Buffer: (A.1) addition of 35 μl pre-heated (56 °C) Elution Buffer to the silica membrane, incubation at room temperature for 3 min, following a centrifugation (11000 g, 1 min); (A.2) addition of a second 35 μl pre-heated (56 °C) Elution Buffer following the same incubation and centrifugation. Alternatively, option B using 70 °C Elution Buffer: (B.1) addition of 50 μl pre-heated (70 °C) Elution Buffer, incubation at room temperature for 3 min and centrifugation at 11000 g for 1 min; (B.2) repetition of the step (B.1) using the first elute.

For the 105 muscle samples, we subsampled approximately 0.5 cm^3^ of muscle from each muscle sample to extract DNA following the standard protocol without any modifications.

### Validation of perch endogenous DNA

DNA concentration of each sample was measured with a Qubit 4 Fluorometer (ThermoFisher Scientific). We used Qubit 1x dsDNA High Sensitivity (HS) and Broad Range (BR) Assay kits to cover the 0.1 to 4000 ng range for both archival bone and contemporary muscle samples.

DNA fragment size distribution was assessed using electrophoresis run on an Agilent 2100 Bioanalyzer system using a DNA 7500 kit. We selected 12 samples (Table S1 & S2) consisting of three individuals from the 1980s and 2000s samples from the two studied populations each to evaluate how storage time affects DNA fragmentation.

To validate the extraction of endogenous perch DNA from the bones, we performed end-point PCR (BIOER thermal cycler) using the QIAGEN Type-it microsatellite PCR kit with two microsatellite primers designed for perch using a subset of all DNA samples that were processed before all other samples. The primers were Pflu4_5 (forward: 5’-TTG ACA TAG CGG TCA AGT CTG T-3’; reverse: 5’-GAT TTG GAT GAC TTG CGT AGG-3’, R. Gross, unpublished) and Pflu4_42 (forward: 5’-CGG ACC AGG TTT CCT ACA GA-3’; reverse: 5’-TGA CTC CAT AAC CCT CCA CA-3’, R. Gross, unpublished). The primers amplified microsatellite loci with size ranges of 115-147 bp (Pflu4_5) and 282-306 bp (Pflu4_42) to evaluate the amplification of fragments with different sizes. Reaction components were added following Type-it Microsatellite PCR (QIAGEN) protocol. The PCR reaction of total volume 15 μl contained 0.3 μl of each four primer of concentration 10 pmol/μl (final concentration in the reaction 0.2 μM), 7.5 μl 2× Type-it Multiplex PCR Master Mix (final concentration 1×), 1.3 μl RNase-free water and 5 μl extracted DNA of concentration < 10 ng/μl. The cycling protocol was as follows: 95°C for 5 min, 35 cycles at 95°C for 30 s, 61°C for 90 s and 72°C for 30 s and a final extension at 60°C for 30 min. PCR products were run on 1% agarose gel stained with ethidium bromide and visualized under a UV light (Bio-Rad GelDoc Go Gel Imaging System).

### Library construction and sequencing

We chose 222 samples (148 bones and 74 muscles, see Table S3) of various DNA concentration (bones: 0.6 – 48.5 ng/μl, muscles: 54-256 ng/μl) and submitted them to the Beijing Genomics Institute (BGI) for library preparation and short read whole genome sequencing using DNBSEQ platform: BGI-SEQ 500 (Goodwin, McPherson, and McCombie 2016; Mak et al. 2017; Zhu et al. 2018). To construct the sequencing libraries of bone DNA samples, the KAPA HyperPrep Kit (Roche) was used as it is designed for highly fragmented DNA in low quantities. The regular DNA Library Prep Kit (Yeasen) was used for the muscle DNA samples. We aimed for 10 × sequencing depth across the genome with read length of 100 bp on pair end mode. No customizations were applied during these steps.

### Bioinformatics

Read quality was assessed using FastQC version 0.11.9 (Andrews, Simon 2019) and all sequences were trimmed with fastp v. 0.23.4 (Chen et al. 2018) applying the parameters: -g - w 12 -r -W 5 -M 25 --trim_ front1 9 --trim_front2 9 --trim_tail1 2 --trim_tail2 2 -l 60 due to the excess of polyG tails (De-Kayne et al. 2021).

Filtered sequence reads of each individual were mapped to the Eurasian perch reference genome (NCBI: GCA_010015445.1) using bowtie2 (Langmead and Salzberg 2012) and SAMtools v. 1.16 (Li et al. 2009). We applied default parameters, with the exception of the modified score minimum threshold (−-score-min L,-0.3,-0.3) and the maximum fragment length for valid paired-end alignments (−X 700). The proportions of mapped reads, coverage and depth were estimated using SAMtools v. 1.16 command flagstat and coverage.

Duplicated reads were identified and marked using Picard v. 2.27.5 MarkDuplicates (Broad Institure 2022). Post mortem DNA damage, such as nucleotides substitution due to deamination (commonly found in aDNA samples, Ginolhac et al. 2011; Sawyer et al. 2012) was assessed using a subset of 36 samples (12 1980s bones, 12 2000s bones and 12 2020s muscles) by Mapdamage2.0 (Jónsson et al. 2013) using default parameters. One of the degradation processes that DNA undergoes over time is cytosine deamination (Briggs et al. 2009), resulting in C to U changes at regular cytosines and C to T changes at 5-methylated cytosines (Briggs et al. 2007). As cytosine deamination mostly occurs at the ends of aDNA fragments (Briggs et al. 2007), we quantified the level of DNA degradation by estimating the percentage of cytosine deamination within 25 bp from the forward read ends.

We carried out Genome Analysis Toolkit (GATK) best practice pipeline v. 4.3.0.0 (Auwera and O’Connor 2020), separately for sequencing data from the nuclear genome and mitochondrial DNA (mtDNA). The HaplotypeCaller subroutine from GATK was applied to the BAM files to generate single-sample GVCF files using the following parameters: -ERC GVCF --minimum-mapping-quality 20 -mbq 13 --G AS_StandardAnnotation. The CombineGVCF tool was then utilized to combine the individuals GVCF files into a single VCF file. Finally, consensus genotypes were called using the GenotypeGVCFs tool.

Initially, 12,775,282 single nucleotide polymorphisms (SNPs) were called using GATK Best Practices pipeline. Subsequently, we employed filtering and retained variants that met the following criteria: (i) mean sequencing depth between 8 and 36 (−-min-meanDP 8 --max-meanDP 36, following (Ozerov et al. 2022); (ii) the consensus quality score of ≥ 30 (−-minQ 30); (iii) minor allele frequency of ≥ 0.05 (MAF, --maf 0.05); (iv) missing in at most 5% of all individuals (−-max-missing 0.95); (v) biallelic sites and not indels (−-max-alleles 2 --min-alleles 2 --remove-indels); and (vi) Hardy-Weinberg equilibrium (HWE) threshold no smaller than 0.01 (−-hwe 0.01).

The procedures and treatments applied to the archive bones during sampling, e.g., bulk processing when removing operculum without following strict disinfection practices, can potentially increase the likelihood of cross-sample contamination. To evaluate this, we calculated observed heterozygosity (*H*_O_) using mitochondrial and nuclear SNPs (hereafter mtDNA *H*_O_ and nDNA *H*_O_) for each sample using PLINK v. 1.90b4.9 --het (Chang et al. 2015). We evaluated whether an individual showed excess or deficiency of heterozygosity by visualizing the distribution of individual *H*_O_ per population and time point.

### Statistical analysis

All data processing and statistical analyses were conducted in R, v. 4.3.3 (R Core Team, 2024). Within bones, we used Analysis of Variance (ANOVA) to test if storage time and elution options affected their DNA yield. Among all samples, we tested if sequencing metrics differed between time points and tissue types (1980s bone, 2000s bone and 2020s muscle) using ANOVA. Data visualization and processing were done using the packages within the tidyverse collection (Wickham et al. 2019).

## RESULTS

### DNA yield and fragmentation

Among all processed bones, the average DNA yield was 307.2 ng (10.6 - 2425.0 ng), while the average DNA yield for muscles was 4972.0 ng (2740 – 12800 ng, Figure 3, Table S1 & S4). Within bones, the average DNA yield was 153.8 ng for the 1980s bones, and 485.5 ng for the 2000s bones. Elution option B using Elution Buffer pre-heated to 70 °C DNA yielded a higher amount of DNA for both the 1980s and 2000s bones than elution option A (Table S1). This indicates that elution option B may demonstrate a higher efficiency in DNA recovery from archival operculum bones. The higher DNA yield was only significant for the 2000s bones (ANOVA, 1980s bones: F(1, 91) = 2.81, P = 0.097; 2000s bones: F(1, 78) = 15.76, P = 1.59e-4). Within bones treated with the same elution option, the 2000s bones yielded a significant higher amount of DNA compared to the 1980s bones (ANOVA, elution option A: F(1, 89) = 12.63, P = 0.61e-4, elution option B: F(1, 80) = 77.25, P = 2.28e-13), suggesting that a negative effect of storage time and DNA yield.

**Figure 3.**
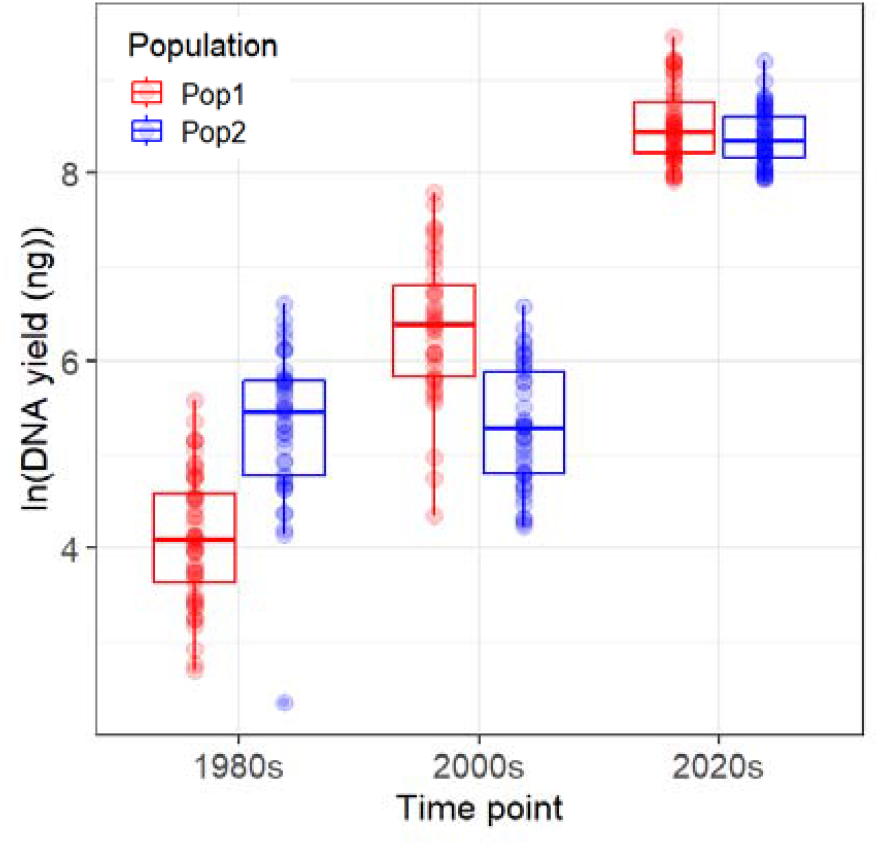
DNA total yield (ng) shown after natural log transformed of the 1980s bones, 2000s bones and 2020s muscle samples from Population 1 (red) and Population 2 (blue). Each point represents an individual DNA sample.

The results of the gel run for the PCR products of 12 bone samples (Figure S2) showed that the amplification of both microsatellite loci was successful for the majority of samples, indicating that the extracted DNA was indeed endogenous perch DNA, with fragments sizes of at least 300 bp. The Bioanalyzer results (Figure S3) showed that most fragments in the 1980s bones were around 300 bp long, while the 2000s bone DNA samples had fragments from 300 bp to up to more than 10000 bp with a peak around 7000 bp.

### Sequencing, mapping and read quality

On average, 70% of reads across all samples were retained after trimming (Figure 4a). Rather unexpectedly, the 1980s bones had the highest proportion of reads retained (mean 71.85%, range 69.90 – 75.77%), while the 2000s bones had the lowest (mean 69.08%, range 63.99 – 73.52%). In general, all samples showed a high proportion of mapped reads (> 95%), except a single outlier individual (< 60%) from the 1980s (Figure 4b). The 2000s bones showed the highest quality with respect to proportion of mapped reads. The level of duplicated reads was low overall (0.7 – 7.8 %), with the 1980s bone samples having the highest proportion of duplicates among the three time points (Figure 4c).

**Figure 4.**
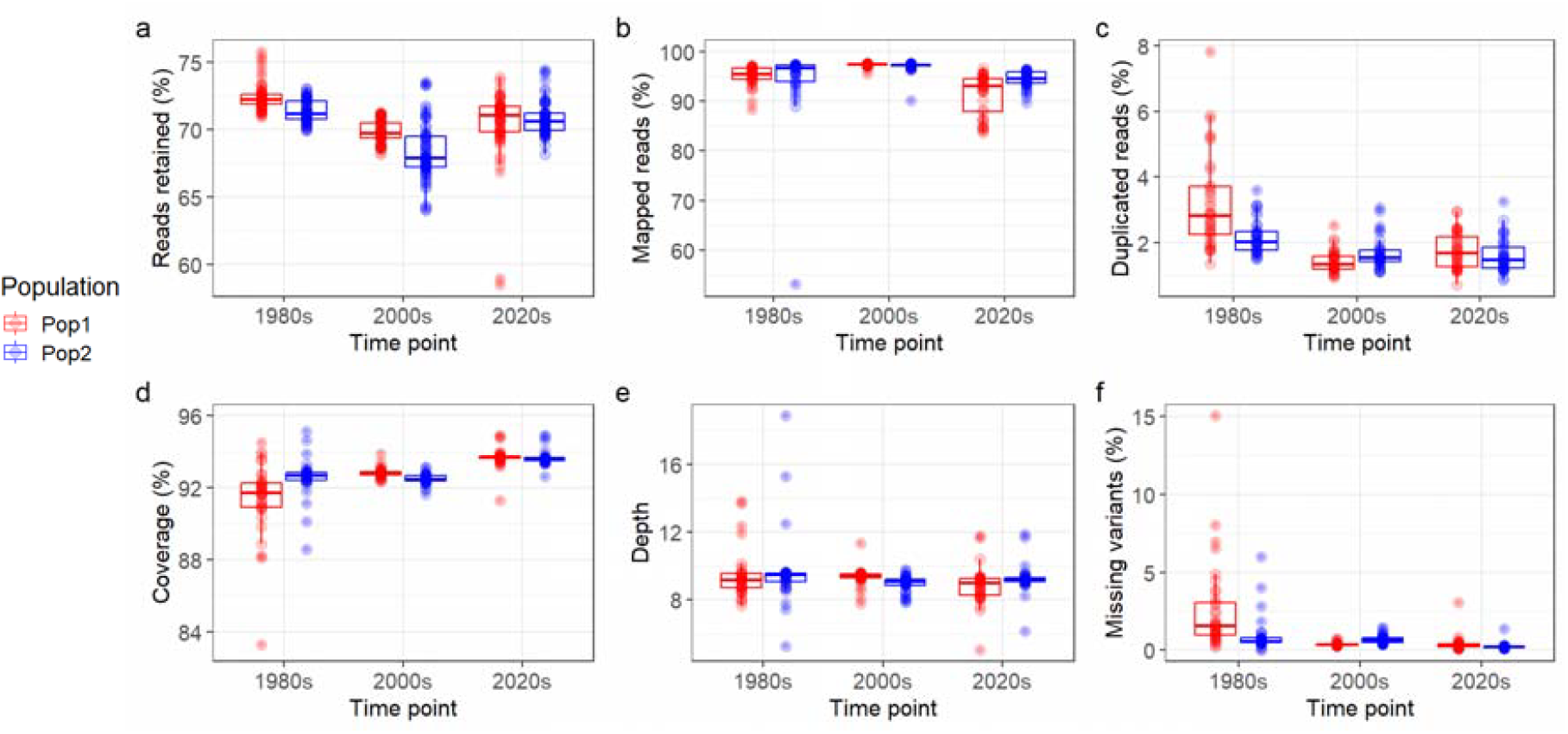
Sequence metrics and mapped reads statistics of the 1980s bone, 2000s bone and 2020s muscle samples from Population 1 (red) and Population 2 (blue): (a) the proportion of reads retained after trimming; (b) the proportion of reads mapped to the reference genome; (c) the percentage of duplicated reads; (d) the percentage of the genome covered by mapped reads; (e) the average depth (the number of times a specific base is covered by sequence reads), and (f) the percentage of missing variants for each individual. Each point represents the genomic data of one individual.

Overall, genome coverage was high (> 90%) across all samples and time points (Figure 4d). Bones from the 1980s had the lowest coverage and most variation among samples, while the contemporary muscle samples showed the highest coverage (>92%). In contrast, sequencing depth was similar between studied samples and time points (Figure 4e), with most samples exceeding 8-fold read depth.

After the filtering, 908,800 SNPs were retained. The percentage of missing variants was low for both bone and muscle samples across all time periods, with a mean missing percentage of 2.37% in the 1980s, 0.52% in the 2000s, and 0.92% in the 2020s. The 1980s bones had the largest variance with some individuals missing more than 5% (Figure 4f).

Using mitochondrial information, a total of six samples exhibited mtDNA *H*_O_ levels above 0.05, including one sample each from the 1980s and 2000s, and four samples from the 2020s, as shown in Figure 5a. Based on nuclear SNP data, a total of eleven individuals (eight samples from the 1980s, one from the 2000s and two from the 2020s) exhibited nDNA *H*_O_ > 0.45 (Figure 5b). However, the excess heterozygosity patterns revealed based on mtDNA *H*_O_ (Figure 5a) and nDNA *H*_O_ (Figure 5b) did not match each other: samples with the highest mtDNA *H*_O_ values were collected in the 2020s, whereas samples with the highest nDNA *H*_O_ estimates were from the 1980s.

**Figure 5.**
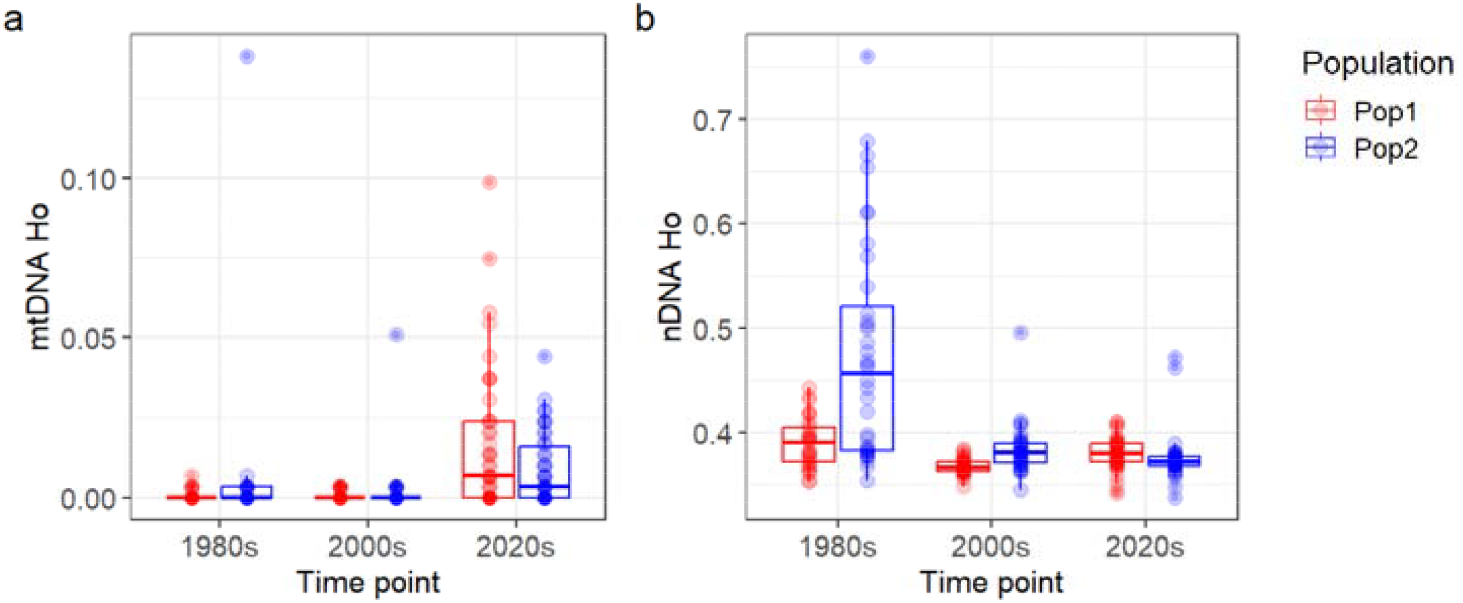
(a) Observed heterozygosity estimated using SNPs from mitochondrial DNA (mtDNA H_O_) and (b) observed heterozygosity estimated using nuclear SNPs (nDNA3 H_O_) for the 1980s bone, 2000s and 2020s samples from Population 1 (red) and Population 2 (blue). Each point represents one individual.

DNA post-mortem damage, as indicated by estimated deamination levels at the forward ends of DNA strands, appeared to be low (<1%) across all samples (Figure 6). The 2000s bones and 2020s muscles were estimated to share similar levels of deamination while the 1980s bones displayed 2-to 4-fold higher deamination levels.

**Figure 6.**
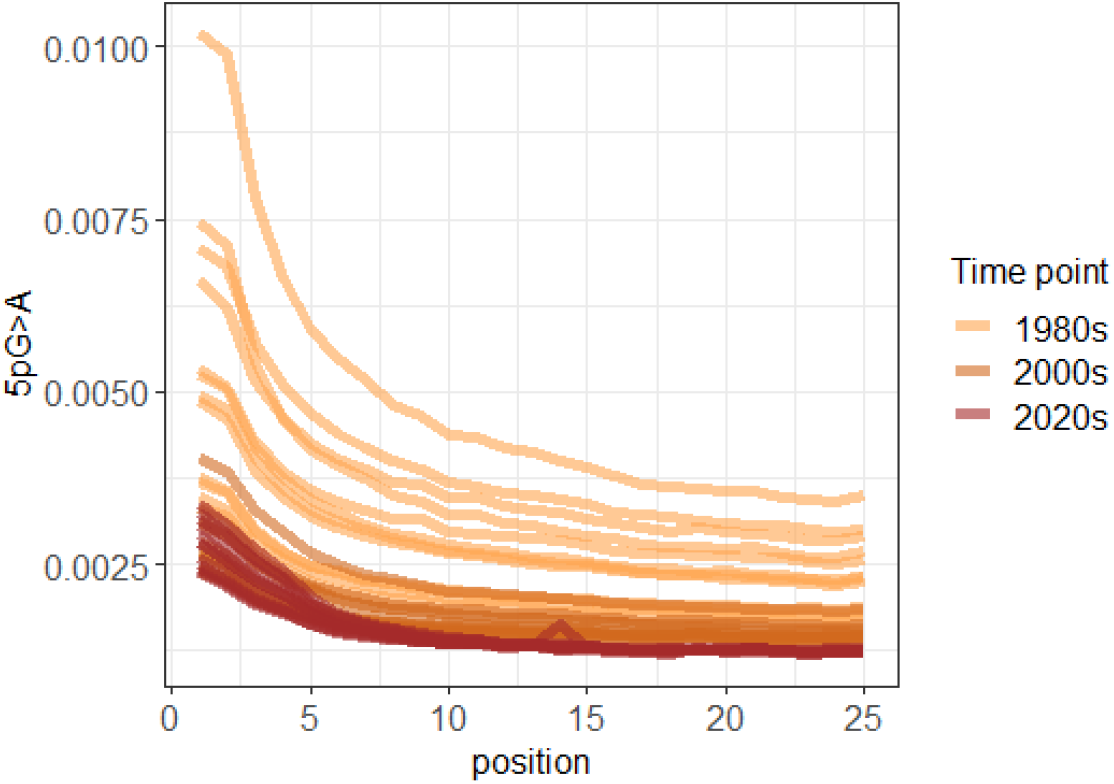
The DNA post-mortem damage patterns shown by 36 samples for the 25 first nucleotides of cytosine to thymine mis-incorporation (equals to G>A) from the forward DNA strand. Each line represents one sample. The darker the colour, the more recent the samples were collected in the order of 1980s bones, 2000s bones and 2020s muscles.

## DISCUSSION

We tested the feasibility of extracting DNA from archival boiled fish operculum bones using a simple protocol. Not only did we succeed, we also compared the quality of subsequent whole genome sequencing data from the boiled fish bones to contemporary muscle tissue. Here, we present our observations made in the process.

As expected, the bone samples yielded significantly less DNA than the muscle samples in terms of both quantity and concentration (Figure 3). However, since bones and muscles are two types of tissues of different properties, the comparison on DNA yield should rather be made between bones of different storage times. The negative effect of storage time (40+ and 20+ years) on DNA yield could be due to the ∼20 additional years of DNA degradation when stored dry at room temperature (Nielsen and Hansen 2008). In addition, DNA extracted from samples with a storage time of 40+ years had much shorter fragments than DNA from samples with storage time of 20+ years. Storage time itself affected both the level of DNA degradation and fragmentation.

To recover the maximal amount of DNA bound to the silica membrane, we experimented with two elution options and observed a slight improvement in elution efficiency when the Elution Buffer was preheated to 70°C. We did not, however, control the volume or bone mass used in the experiment, so we cannot definitely conclude that preheating Elution Buffer to 70°C improves the DNA yield. Additionally, some DNA extraction protocols, such as the Monarch® Spin gDNA Extraction Kit (NEB #T3010S/L), recommend against preheating the elution buffer to temperatures above 60°C, as hot elution may result in partial and irreversible denaturation of the eluted genomic DNA, depending on the salt conditions. Readers should exercise caution when determining the elution step.

More importantly, based on the general patterns in sequencing metrics, the bones from the 1980s, 2000s, and muscles from the 2020s all exhibited high sequence data quality (Figure 4). With the only exception of genome coverage (Figure 4d), the bones generally displayed equal or higher quality levels compared to the muscles, suggesting that neither the type of tissue, storage time, nor boiling treatment had a major negative impact on the quality of the sequencing data. Notably, boiled archival operculum bones can yield whole genome sequencing data of quality comparable to that of fresh muscle samples.

We evaluated the level of cross-contamination by estimating observed heterozygosity of each sample and were left with contradicting patterns based on mtDNA (Figure 5a) and nuclear SNPs (Figure 5b). However, given that the nuclear SNP dataset is much larger compared to SNPs on mtDNA, it is likely that the observed heterozygosity estimates based on nuclear data provide more robust and reliable information about potential contamination than mtDNA H_O._ Thus, it is likely that the highest level of cross-individual contamination was observed in the population 2 samples from the 1980s. This result also highlights batch-effect in relation to cross-contamination (during sampling) when studying archival samples. As the bones were originally collected for age and growth determination rather than DNA extraction (Thoresson 1996), their sampling process lacked standard molecular analysis precautions and decontamination steps, such as changing gloves and sterilizing boiling-water chambers in between samples. In comparison, the muscle sampling process followed a more rigorous decontamination routine. We also observe slightly increased level of DNA degradation in the 1980s bones in terms of post-mortem damage (Figure 6). Nevertheless, we only detected a few low quality archival DNA samples and those can be easily removed by filtering for downstream genomic analysis.

During recent years, first successful attempts have been published to obtain whole genome sequencing data using archival otolith and scales, demonstrating that fish bones are a good source of DNA (Ferrari et al. 2021). Furthermore, recent studies have provided much-needed temporal perspective to investigate the association between concurrent environmental changes and evolutionary changes in wild fish populations (Caccavo et al. 2024; Pinsky et al. 2021). As a next step, we are currently investigating the WGS data acquired from the two populations to identify selection signatures of rising water temperatures, aiming to provide evolutionary insights on how fish genomes evolve and change in the context of climate change (Niu et al. in prep.).

We hope our demonstration of a simple and efficient DNA isolation protocol on alternative bony samples motivates further utilization of archives in fisheries institutions and museums. Further optimizing the extraction protocol from fish operculum bones would require additional investigation into factors such as bone thickness and size, which may influence DNA yield and quality. Understanding these relationships is important for developing effective protocols for genomic analyses using operculum bones. Accordingly, the field of fish genomics would benefit largely from a broader and more systematic evaluation of methods and protocols for DNA extraction from more alternative types of archival bony material and the subsequent library preparation and sequencing practices. Our findings highlight that although the sampling (e.g., the boiling) and storage were suboptimal for molecular analyses, the utility of archival fish operculum bones for evolutionary studies is comparable to fresh muscle samples. Overall, the integration of genomic, archaeological and ecological approaches offers the potential to address a range of scientific and management questions that cannot be fully achieved by either discipline in isolation.

## Authors’ contributions

J.N.: conceptualization, data curation, formal analysis, investigation, methodology, visualization, writing—original draft, writing—review and editing;

A.V.: conceptualization, investigation, methodology, funding acquisition, supervision, writing—original draft, writing—review and editing

M. E. L.: data curation, formal analysis, investigation, methodology, supervision, writing— original draft, writing—review and editing

L. P.: data curation, investigation, methodology, supervision, writing—review and editing

M. H.: conceptualization, data curation, writing—review and editing

A. G.: conceptualization, data curation, funding acquisition, supervision, writing—original draft, review and editing

## Acknowledgements

We are grateful for the guidance and help received from Prune Leroy, Lauren Davis, Javier Edo Varg when developing the extraction protocol and from Katarina Ihrmark when conducting the Bioanalyzer analyses. We thank past and current staff for fish tissue sampling and archival sample maintenance in Öregrund, Sweden. The computations and data handling were enabled by resources in project NAISS2023-5-114, NAISS2024-22-446 and NAISS2024-6-93 provided by the Swedish National Infrastructure for Computing (SNIC) at UPPMAX. This study was funded by FORMAS (Grant 2019-00928 to AG), the Oscar and Lili Lamm Foundation (to AG), Estonian Research Council (Grant PRG852 to AV) and SLU Quantitative Fish- and Fisheries Ecology.

