## Supplementary materials for "DNA extracted from boiled archival fish bones yields high quality whole genome sequencing data"

### Although we selected bones from fish of as similar sizes as possible, there is a wide range of fish sizes (Figure S1).

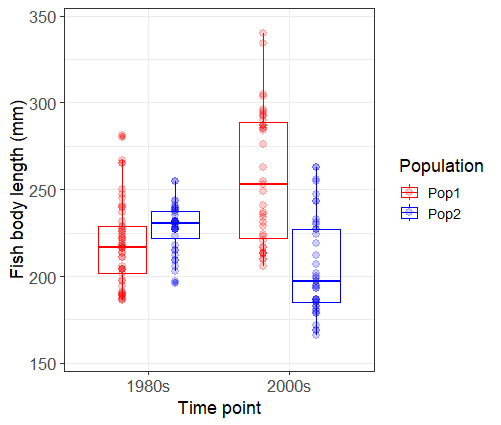

Figure S1. Size distributions of the fish from which the operculum bones were selected in this study. Each point represents an individual fish, red for Population 1 and blue for Population 2.

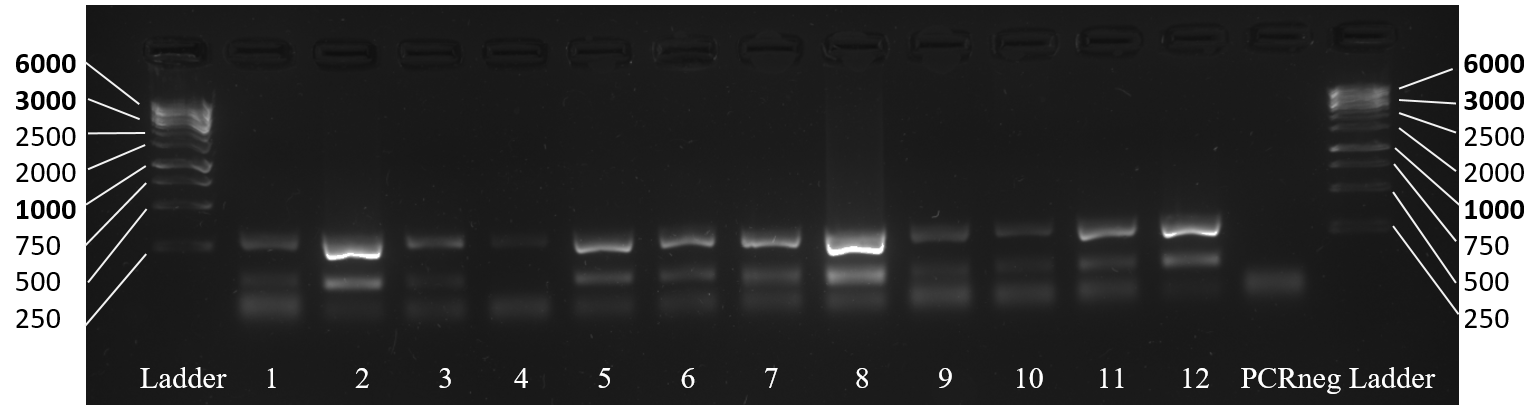

Figure S2. Multiplex PCR of P. fluviatilis microsatellite loci (Pflu4_5, 115-147 bp; Pflu4_42, 282-306 bp) using 12 bones samples visualized in 1% ethidium bromide stained agarose gel. The columns indicated as “Ladder” were Thermo Scientific GeneRuler 1 kb DNA Ladders, used to indicate the fragment size (bp). The lowest band shown in all samples including the PCR negative control (PCRneg) was the amplification of the primers (~20 bp). Besides the primer band, all except bone sample 4 showed two bands, suggesting the amplification of both loci.

We assessed the distribution of DNA fragment size of 12 DNA samples (six 1980s bones and six 2000s bones, Figure S2). We show one example each for the 1980s and 2000s in Figure S3. Bioanalyzer result of all 12 samples (Table S1) are shown in the supplementary file bioanalyzer.12sample.kit7500.pdf.

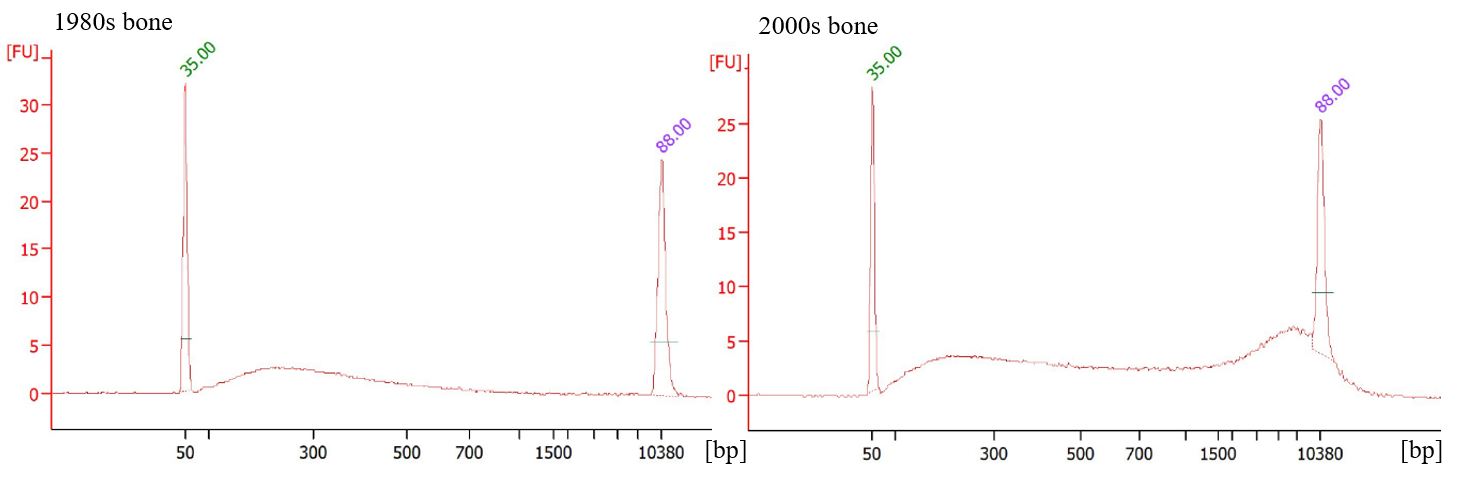

Figure S3. One example each of DNA fragment size distribution from the 1980s bones (left) and 2000s bones (right) analysed by the DNA 7500 kit for 2100 Bioanalyzer Systems. From low to high numbers along the x axis, each tick representing the 50, 100, 300, 500, 700, 1000, 1500, 2000, 3000, 5000, 7000 and 10380 bp marker of the ladder. The 1980s bone shows only one bump just below 300 bp, indicating that most fragments were around 300 bp long. The 2000s bone shows a distribution curve that elevates from below 300 bp, slightly and slowly decreases after, and peaks around 7000 bp. This shows that a significant amount of fragments ranging 300 - 10000 bp were present in the 2000s sample.

Table S2. The key to the 12 bone samples presented in bioanalyzer.12sample.kit7500.pdf. They were also marked in Table S1.

| Sample | Name | Time point |
| --- | --- | --- |
| 1 | Yao_sample_BT1 | 1980s |
| 2 | Yao_sample_BT2 | 1980s |
| 3 | Yao_sample_BT3 | 1980s |
| 4 | Yao_sample_BT4 | 2000s |
| 5 | Yao_sample_BT5 | 2000s |
| 6 | Yao_sample_BT6 | 2000s |
| 7 | Yao_sample_FM1 | 1980s |
| 8 | Yao_sample_FM2 | 1980s |
| 9 | Yao_sample_FM3 | 1980s |
| 10 | Yao_sample_FM6 | 2000s |
| 11 | Yao_sample_FM4 | 2000s |
| 12 | Yao_sample_FM5 | 2000s |
